# Avian thermoregulation in the heat: phylogenetic variation among avian orders in evaporative cooling capacity and heat tolerance

**DOI:** 10.1101/211730

**Authors:** Ben Smit, Maxine C. Whitfield, William A. Talbot, Alexander R. Gerson, Andrew E. McKechnie, Blair O. Wolf

## Abstract

Little is known about the phylogenetic variation of avian evaporative cooling efficiency and heat tolerance in hot environments. We quantified thermoregulatory responses to high air temperature (*T*_a_) in ~100-g representatives of three orders: African cuckoo (*Cuculus gularis*, Cuculiformes), lilac-breasted roller (*Coracias caudatus*, Coraciiformes), and Burchell’s starling (*Lamprotornis australis*, Passeriformes). All three species initiated respiratory mechanisms to increase evaporative heat dissipation when body temperature (*T*_b_) approached 41.5°C in response to increasing *T*_a_, with gular flutter observed in cuckoos and panting in rollers and starlings. Resting metabolic rate (RMR) and evaporative water loss (EWL) increased by quantitatively similar magnitudes in all three species, although maximum rates of EWL were proportionately lower in starlings. Evaporative cooling efficiency [defined as the ratio of evaporative heat loss (EHL) to metabolic heat production (MHP)] generally remained below 2.0 in cuckoos and starlings, but reached a maximum of ~3.5 in rollers. The high value for rollers reveals a very efficient evaporative cooling mechanism, and is similar to EHL/MHP maxima for similarly sized columbids which very effectively dissipate heat via cutaneous evaporation. This unexpected phylogenetic variation among the orders tested in the physiological mechanisms of heat dissipation is an important step toward determining the evolution of heat tolerance traits in desert birds.

**Summary statement:** We show that avian evaporative cooling efficiency and heat tolerance display substantial taxonomic variation that are, unexpectedly, not systematically related to the use of panting versus gular flutter processes.

## Introduction

Evaporative heat loss (EHL) is the only mechanism whereby birds can maintain body temperature (*T*_b_) below lethal limits in hot environments where environmental temperature exceeds *T*_b_. Rapid increases in evaporative water loss (EWL) are a ubiquitous avian response to such conditions (Bartholomew and Cade, 1963; Dawson and Bartholomew, 1968; Smith et al., 2015; Whitfield et al., 2015). Water requirements for thermoregulation have thus shaped the ecology and evolution of birds living in hot environments, and provide the basis for important trade-offs between dehydration and hyperthermia avoidance in arid-zone species (Smit et al., 2013; Tieleman and Williams, 2000; Tieleman and Williams, 2002; Tieleman et al., 2003).

Avian heat tolerance and evaporative cooling efficiency (quantified as the maximum ratio of heat dissipated evaporatively, EHL, to that generated metabolically, MHP) appears to vary substantially among orders (Lasiewski and Seymour, 1972; Smith et al., 2015), within orders (McKechnie et al., 2016a; McKechnie et al., 2017), and even within species (Noakes et al., 2016). For example, recent studies have shown that the efficiency of evaporative cooling is generally very high (EHL/MHP > 2.0) in members of the orders Columbiformes and Caprimulgiformes (McKechnie et al., 2016a; O’Connor et al., 2017; Smith et al., 2015; Talbot et al., 2017), but less so in Passeriformes, Pteroclidiformes, Galliformes and Strigiformes, where efficiency rarely exceeds 2.0 (Bartholomew et al., 1968; McKechnie et al., 2016b; McKechnie et al., 2017; Smith et al., 2015; Smith et al., 2017; Whitfield et al., 2015). Narrow phylogenetic sampling means we still have an incomplete understanding of the diversity of avian EHL mechanisms and their functional significance for heat tolerance.

Body mass represents one of the most prominent sources of variation in heat tolerance and the efficiency of EHL (McKechnie and Wolf, 2010), but even species of similar size can still show substantial variation. For example, Lasiewski and Seymore (1972) showed that four similarly-sized species from different orders (*Ploceus cucullatus*, Passeriformes; *Excalfactoria chinensis*, Galliformes; *Scardafella inca*, Columbiformes; *Phalaenoptilus nuttallii*, Caprimulgiformes) show substantial variation in terms of the magnitude of elevations in *T*_b_, metabolism, EWL, and EHL/MHP during heat exposure. Recent studies have suggested that some of the mechanisms underlying this variation may include differential reliance on respiratory versus cutaneous pathways of evaporative cooling (McKechnie et al., 2016a), and morphological variation (e.g. bill size) (Danner et al., 2016; Tattersall et al., 2009; van de Ven et al., 2016). Species relying on cutaneous evaporative pathways, such as Columbiformes, are thought to show negligible metabolic heat production associated with evaporative cooling demands (McKechnie et al., 2016a). In contrast, species relying on respiratory evaporative cooling (especially panting) may incur greater metabolic costs in their efforts to dissipate heat (Bartholomew et al., 1968; Lasiewski and Seymour, 1972). Gular flutter (gaping while pulsating the hyoid bone to ventilate the buccal cavity) has been argued to require less energy than panting (gaping while using rapid movements of the thorax and abdominal cavities) (Bartholomew et al., 1962; Dawson, 1958; Lasiewski and Bartholomew, 1966); the gular flutter process may thus enhance efficiency of EHL. The use of gular flutter *versus* panting further seems to vary among taxa (Calder and Schmidt-Nielsen, 1967), but the phylogenetic distribution of these evaporative cooling processes remains poorly understood.

Variation in the rate and efficiency of evaporative cooling may reflect physiological costs, such as risk of dehydration and hyperthermia (McKechnie and Wolf, 2010), but may also reflect behavioural [e.g. reduced foraging efficiency correlated with heat dissipation efforts (du Plessis et al., 2012)] and ecological costs [e.g. reliance of surface water sources (Smit et al., 2016)]. Thus far, evaporative cooling efficiency and heat tolerance are best-studied in the Passeriformes and Columbiformes, and data from more under-represented orders are needed to further our understanding of the evolutionary drivers and mechanisms involved in thermoregulation in the heat.

Here we present data on thermoregulation under very hot conditions in three 100-g species from three orders: Cuculiformes, Coraciiformes and Passeriformes. The Cuculiformes and Coraciiformes remain largely unstudied in terms of thermoregulation at high environmental temperatures; both represent diverse taxa that include many species occupying hot, arid regions. To the best of our knowledge, the only thermoregulatory data from a cuculiform under hot conditions is for the roadrunner, *Geococcyx californianus* (Calder and Schmidt-Nielsen, 1967). In the latter study, high chamber humidity may have reduced the efficiency of EHL at air temperatures (*T*_a_) below *T*_b_ (Gerson et al., 2014). Recent advances in carbon dioxide and water analysers have allowed us to maintain low humidity levels during testing by using very high flow rates, thus avoiding the complications associated with high humidity that plagued early studies (Gerson et al., 2014; Lasiewski et al., 1966; Smith et al., 2015; Whitfield et al., 2015). We further describe the mechanism of respiratory EHL (panting or gular flutter) used by these three species, and predict that, compared to panting, the use of gular flutter is strongly associated with improved evaporative cooling efficiency and heat tolerance. In addition, our inclusion of a large passerine (>100 g) allows for more rigorous analyses of the scaling of traits related to heat tolerance and evaporative cooling in this speciose taxon (McKechnie et al., 2017).

## Methods

### Study species and study sites

We measured resting metabolic rate (RMR), evaporative water loss (EWL), and *T*_b_ in species representing three avian orders in the southern Kalahari Desert in the Northern Cape province of South Africa, an arid area with a mean annual rainfall of ~200 mm and summer daily maximum *T*_a_ ranging from ~20-43°C (Whitfield et al., 2015). We followed the methodology of Whitfield et al. (2015) to quantify thermoregulatory responses to high *T*_a_ in a field laboratory from January to April 2012 at Wildsgenot Game Ranch (27°04′S, 21°23′E), and from January to March 2013 at Leeupan Game Ranch (26°58′S, 21°50′E).

We trapped six African cuckoos (*Cuculus gularis* Stephens; order Cuculiformes, hereafter cuckoos) with a mean ± s.d. body mass (*M*_b_) of 109.6 ± 5.6 g (range 99.0 - 116.0 g) at Leeupan Game Ranch during early 2013 (austral summer) using mist nets. We trapped 10 lilac-breasted rollers (*Coracias caudatus* Linnaeus; Order Coraciiformes, hereafter rollers) with a *M*_b_ of 95.4 ± 8.5 g (78.2 - 110.35 g); two individuals were trapped in February 2012 at Wildsgenot Game Ranch, and the remaining individuals were trapped at Leeupan Game Ranch between January 2013 and March 2013. We trapped seven Burchell’s starlings (*Lamprotornis australis* Smith; order Passeriformes, hereafter starlings) with a *M*_b_ of 109.1 ± 9.3 g (85.4 - 116.9 g) using mist nets and/or flap traps baited with tenebrionid beetle larvae on Wildsgenot Game Ranch in February 2012, and Leeupan Game Ranch from December 2013 to February 2013. Measurements were carried out on the same day of capture. If birds were not tested within the first three hours after being trapped, they were housed in cages constructed of shade-cloth, and were provided with tenebrionid beetle larvae and water ad libitum, until physiological measurements commenced.

Birds were held in respirometry chambers for 2-3 hr, a period that typically limited *M*_b_ loss to <5% of initial *M*_b_ (mean *M*_b_ loss during measurements was 4.0 ± 1.8% of initial values) and time in captivity did not exceed 24 hr, after which birds were released at the site of capture following Whitfield et al. (2015). All experimental procedures were approved by the Animal Ethics Committee of the University of Pretoria (protocol EC071-11) and the Institutional Animal Care and Use Committee of the University of New Mexico (12-1005370-MCC). A permit to trap the birds was issued by the Northern Cape Department of Environmental Affairs (ODB 008/2013).

### Gas exchange and temperature measurements

We used the same experimental set-up as described by Whitfield et al. (2015) to obtain gas exchange, *T*_a_ and *T*_b_ measurements. We measured *T*_b_ using calibrated temperature-sensitive passive integrated transponder (PIT) tags (Biomark, Boise, ID, USA) which were injected intraperitoneally into the abdominal cavity of each bird shortly after capture. We monitored *T*_b_ throughout gas exchange measurements, using a reader and transceiver system (model FS2001, Biomark, Boise, ID, USA). We obtained carbon dioxide production (V_CO_2__) and EWL measurements over the *T*_a_ range of 25-56°C (depending on the species), also using the same experimental setup as described by Whitfield et al. (2015). All three species were placed individually in 9-L plastic chambers, and stood on a platform of plastic mesh 10 cm above a 1-cm layer of mineral oil to trap excreta. We used flow rates between 9 and 30 L min^−1^ for cuckoos; 9 and 70 L min^−1^ for rollers, and 10 and 55 L min^−1^ for starlings, depending on the experimental *T*_a_, in order to keep chamber humidity below 5 ppt. As was the case in previous studies using the same methodology (Smith et al. 2015, Whitfield et al. 2015, McKechnie et al. 2016a, McKechnie et al. 2016b), birds remained calmer at very high *T*_a_ when we increased flow rate (and hence decreased humidity).

### Experimental protocol

Following the protocol described by Whitfield et al. (2015), we exposed birds to progressively higher *T*_a_, with increments of approximately 5°C between 25 and 40°C, and 2°C increments between 40°C and the maximum *T*_a_ (ranging from 48 to 56°C depending on the species). Tests were conducted during the day, as this was the active phase of all three species. Birds spent between 10 and 30 min at each *T*_a_ value. We continually monitored birds during measurements using a live video feed and an infrared light source (Whitfield et al., 2015). We recorded behavioural responses of birds within the respirometry chambers every two minutes; an activity state score of the individual (e.g. calm, turning around or jumping) was recorded, as well as respiratory EHL mechanisms, including panting (defined as gaping) and gular flutter (defined as an obvious pulsation of hyoid bone in the gular area while gaping). For each individual, we recorded the air temperature and body temperature at which heat dissipation behaviour was initiated.

We followed Whitfield et al. (2015) in terminating test runs when a bird a) exhibited prolonged escape behaviour such as agitated jumping, pecking and/or wing flapping), b) showed signs of a loss of coordination or balance accompanied by *T*_b_ > 44°C, or c) exhibited a decrease in EWL and RMR accompanied by an uncontrolled increase in *T*_b_. In the last instance, a bird was considered to have reached its upper limit of heat tolerance, and the *T*_a_ associated with the onset of these signs of heat stress was considered the thermal endpoint for that individual. When a bird reached any the conditions described above, we removed the individual from the chamber by hand, gently rubbed a cotton pad soaked in ethanol onto its body, and held it in front of an air-conditioner producing chilled air in order to facilitate rapid heat loss (Whitfield et al., 2015).

### Data analyses

Data were analysed following Whitfield et al.( 2015). We calculated physiological estimates by extracting, EWL and *T*_b_ as the lowest 5 min mean at each *T*_a_ using Expedata (Sable Systems, Las Vegas NV, USA). We present whole-animal values, although we also calculated the slope of mass-specific EWL vs. *T*_a_ to compare our values with the allometric equation presented by McKechnie and Wolf (2010). We followed McKechnie et al. (2017) by converting rates of EWL to evaporative heat loss (W) assuming a latent heat of vaporisation of water of 2.406 J mg^−1^ at 40°C (Tracy et al., 2010). Since none of our study species have crops we assumed all birds were post-absorptive at the time of measurements, but we were unable to confirm this. Measurements typically took place more than 1 hr after capture, and food (mealworm larvae) was only offered to individuals not tested within 3 hrs of capture. We therefore assumed a respiratory exchange ratio of 0.85, representative of a mix of carbohydrate and lipid metabolism in post-absorptive birds (Walsberg and Wolf 1995), and converted rates of V_CO_2__ to metabolic rate (Watt, W) using 24.4 J mL^−1^ CO_2_ (Withers 1992).

We used segmented linear regression models fitted in the *R* package *segmented* (Muggeo, 2009) to estimate inflection *T*_a_ in the physiological data. Although our use of these models violates assumptions of independent samples, we used segmented models purely to aid us in identifying inflection *T*_a_ values in physiological variables. We subsequently used linear mixed effects models that included individual identity as a random factor in the *R* package *nlme* (Pinheiro et al., 2009), to obtain estimates of physiological variables as a function of *T*_a_, using subsets of the data above respective inflection *T*_a_. We contrasted the slopes of physiological parameters obtained from these mixed effects models among species, by including data previously collected from laughing doves (*Stigmatopelia senegalensis*, 89.4 ± 13.0 g, hereafter doves) studied at the same time and site as our species (McKechnie et al., 2016a), and freckled nightjars (*Caprimulgus tristigma*, 64.7 ± 6.3 g, hereafter nightjars) studied from a desert site in Namaqualand, South Africa (O’Connor et al., 2017). Thermoregulatory data in the doves and nightjars were collected in the same manner as described here, although we recalculated physiological estimates of V_CO_2__, EWL and *T*_b_ in the dove, as the lowest 5 min mean at each *T*_a_, to facilitate comparison with our study species and the nightjar. Our small sample size of species prevented us from conducting more rigorous statistical analyses that test phylogenetic inertia and, if necessary, account for phylogenetic relatedness. We calculated the upper and lower 95% confidence limits (CL) of each coefficient, and considered species to differ quantitatively in the parameters when there was no overlap in the 95% CL.

## Results

### Body temperature

The *T*_b_ measured at *T*_a_s below the respective *T*_uc_ of each species varied by 0.7°C, from a mean of 40.0 ± 0.6°C in cuckoos to 40.7 ± 0.8°C in rollers (Table 1). The inflection of *T*_a_ above which *T*_b_ increased was < 37°C in all three species (Table 1), with *T*_b_ increasing linearly and significantly above the inflection *T*_a_s (cuckoo, t_1,26_=14.275, p<0.001; rollers, t_1,56_= 12.10, p<0.001; starlings, t_1,28_=7.71, p<0.001) (Fig. 1). The maximum *T*_a_ reached was 50°C for both cuckoos and starlings, and 56°C for rollers. Any further increases in *T*_a_ resulted in these individuals becoming severely agitated. Average maximum *T*_b_ ranged between 42.8° and 44.0°C in the three species at the highest *T*_a_s tested. In two calm cuckoos, *T*_b_ exceeded 44°C at *T*_a_ = 46° and 48°C, respectively, whereupon they lost their ability to right themselves and the experiment was terminated immediately. With the exception of *T*_b_ > 44°C in one roller individual (with no ill-effects), *T*_b_ remained < 44°C in calm rollers and starlings at the maximum *T*_a_s. At the highest *T*_a_ shared among the three species (46°C), rollers maintained a lower *T*_b_ than cuckoos and starlings (Table 1). The mean *T*_a_ at which birds initiated respiratory EHL mechanisms varied from 35.9°C in starlings to ~41°C in cuckoos and rollers, typically as *T*_b_ approached 41.5°C in all three species (Table 1). Our observations of the cuckoos revealed that gaping was always accompanied by gular flutter, which was visibly evident from the pulsating throat and moving hyoid apparatus. Rollers and starlings panted while gaping (the gape is wide in rollers) and we did not observe gular flutter (hyoid movement) in either species.

**Figure 1:**
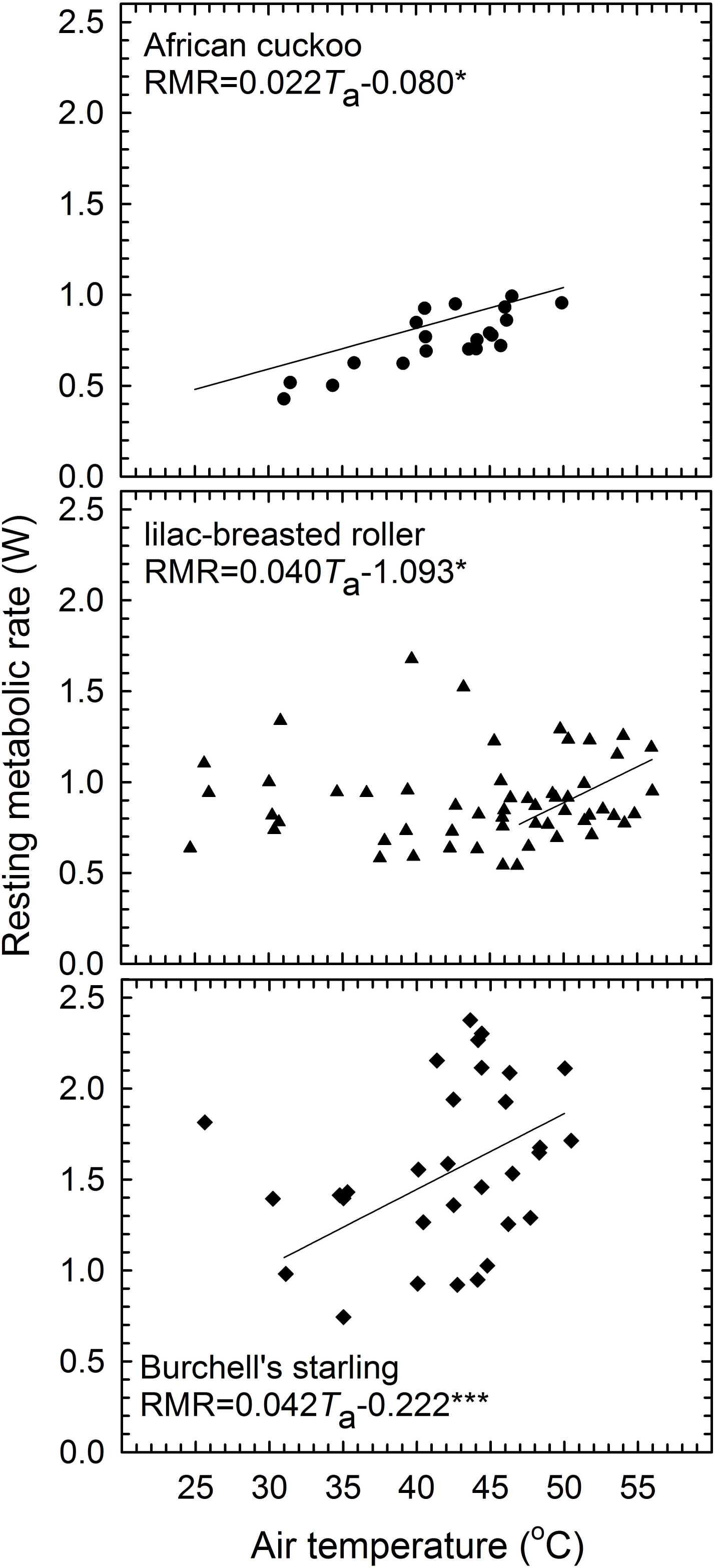
Body temperature (*T*_b_) in African cuckoos (*Cuculus gularis*; n=6), lilac-breasted rollers (*Coracias caudatus*; n=10), and Burchell’s starlings (*Lamprotornis australis*; n=7) as a result of air temperature (*T*_a_). The trendlines represent relationships between *T*_b_ and *T*_a_ above an upper inflection *T*_a_ (see methods). The slopes and intercepts were calculated using linear mixed-effects models. Significant relationships are represented by asterisks: * = p < 0.05, ** = p < 0.01, *** = p < 0.001.

**Table 1:**
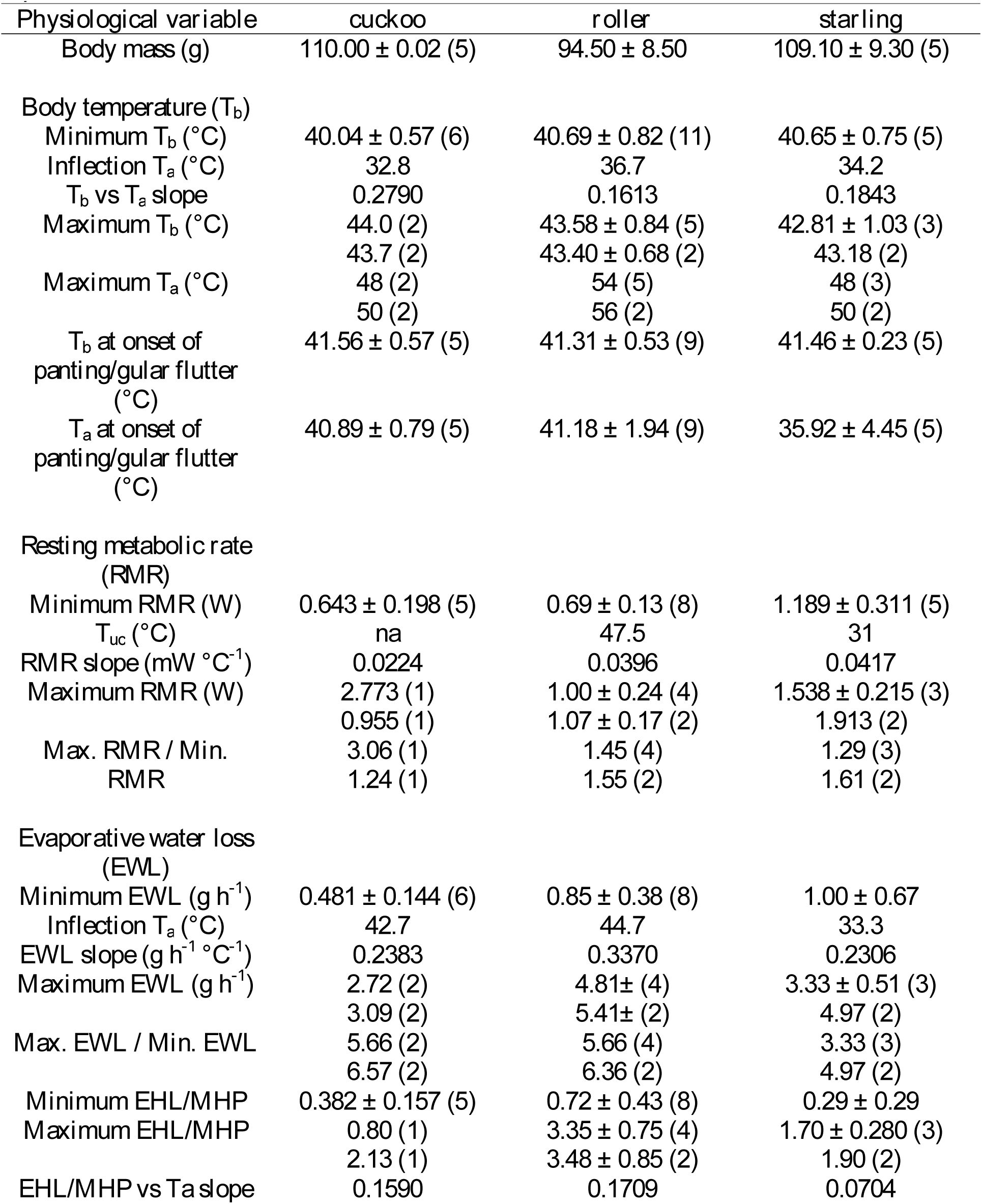
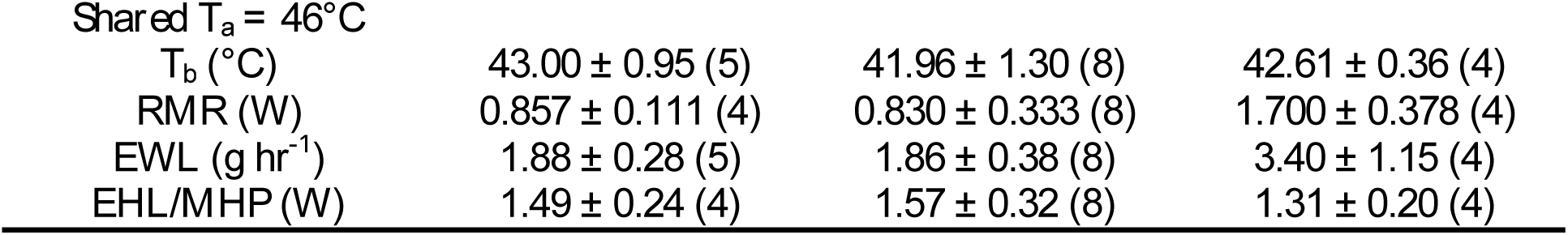
Physiological variables (mean ± s.d., n) related to thermoregulation at the air temperature (*T*_a_) range we tested African cuckoo (*Cuculus gularis*; cuckoo), lilac-breasted roller (*Coracias caudatus;* roller), and Burchell’s starling (*Lamprotornis australis;* starling). Minimum values represent lowest value for each individual at *T*_a_ < 35°C, or the upper critical limit of thermoneutrality (*T*_uc_) for each species. Maximum values where obtained at the highest *T*_a_ tested; when n<3, we also show average values at the second highest *T*_a_ tested. We further indicate mean values for the highest *T*_a_ (n≥3), i.e. *T*_a_ = 46°C, shared by the three species.

| Physiological variable | cuckoo | roller | starling |
| --- | --- | --- | --- |
| Body mass (g) | 110.00 ± 0.02 (5) | 94.50 ± 8.50 | 109.10 ± 9.30 (5) |
| Body temperature ($T_b$) | | | |
| Minimum $T_b$ ($^\circ\text{C}$) | 40.04 ± 0.57 (6) | 40.69 ± 0.82 (11) | 40.65 ± 0.75 (5) |
| Inflection $T_a$ ($^\circ\text{C}$) | 32.8 | 36.7 | 34.2 |
| $T_b$ vs $T_a$ slope | 0.2790 | 0.1613 | 0.1843 |
| Maximum $T_b$ ($^\circ\text{C}$) | 44.0 (2) | 43.58 ± 0.84 (5) | 42.81 ± 1.03 (3) |
| | 43.7 (2) | 43.40 ± 0.68 (2) | 43.18 (2) |
| Maximum $T_a$ ($^\circ\text{C}$) | 48 (2) | 54 (5) | 48 (3) |
|  | 50 (2) | 56 (2) | 50 (2) |
| $T_b$ at onset of panting/gular flutter ($^\circ\text{C}$) | 41.56 ± 0.57 (5) | 41.31 ± 0.53 (9) | 41.46 ± 0.23 (5) |
| $T_a$ at onset of panting/gular flutter ($^\circ\text{C}$) | 40.89 ± 0.79 (5) | 41.18 ± 1.94 (9) | 35.92 ± 4.45 (5) |
| Resting metabolic rate (RMR) |  |  |  |
| Minimum RMR (W) | 0.643 ± 0.198 (5) | 0.69 ± 0.13 (8) | 1.189 ± 0.311 (5) |
| $T_{uc}$ ($^\circ\text{C}$) | na | 47.5 | 31 |
| RMR slope (mW $^\circ\text{C}^{-1}$) | 0.0224 | 0.0396 | 0.0417 |
| Maximum RMR (W) | 2.773 (1) | 1.00 ± 0.24 (4) | 1.538 ± 0.215 (3) |
| | 0.955 (1) | 1.07 ± 0.17 (2) | 1.913 (2) |
| Max. RMR / Min. RMR | 3.06 (1) | 1.45 (4) | 1.29 (3) |
|  | 1.24 (1) | 1.55 (2) | 1.61 (2) |
| Evaporative water loss (EWL) |  |  |  |
| Minimum EWL (g h$^{-1}$) | 0.481 ± 0.144 (6) | 0.85 ± 0.38 (8) | 1.00 ± 0.67 |
| Inflection $T_a$ ($^\circ\text{C}$) | 42.7 | 44.7 | 33.3 |
| EWL slope (g h$^{-1}$ $^\circ\text{C}^{-1}$) | 0.2383 | 0.3370 | 0.2306 |
| Maximum EWL (g h$^{-1}$) | 2.72 (2) | 4.81 ± (4) | 3.33 ± 0.51 (3) |
| | 3.09 (2) | 5.41 ± (2) | 4.97 (2) |
| Max. EWL / Min. EWL | 5.66 (2) | 5.66 (4) | 3.33 (3) |
|  | 6.57 (2) | 6.36 (2) | 4.97 (2) |
| Minimum EHL/MHP | 0.382 ± 0.157 (5) | 0.72 ± 0.43 (8) | 0.29 ± 0.29 |
| Maximum EHL/MHP | 0.80 (1) | 3.35 ± 0.75 (4) | 1.70 ± 0.280 (3) |
| | 2.13 (1) | 3.48 ± 0.85 (2) | 1.90 (2) |
| EHL/MHP vs $T_a$ slope | 0.1590 | 0.1709 | 0.0704 |
| *Shared $T_a = 46^\circ\text{C}$* |  |  |  |
| $T_b$ ($^\circ\text{C}$) | $43.00 \pm 0.95$ (5) | $41.96 \pm 1.30$ (8) | $42.61 \pm 0.36$ (4) |
| RMR (W) | $0.857 \pm 0.111$ (4) | $0.830 \pm 0.333$ (8) | $1.700 \pm 0.378$ (4) |
| EWL ($\text{g hr}^{-1}$) | $1.88 \pm 0.28$ (5) | $1.86 \pm 0.38$ (8) | $3.40 \pm 1.15$ (4) |
| EHL/MHP (W) | $1.49 \pm 0.24$ (4) | $1.57 \pm 0.32$ (8) | $1.31 \pm 0.20$ (4) |

### Resting metabolic rate

One cuckoo showed abnormally high RMR values (ranging from 1.03 W at *T*_a_ = 30°C, to 5.20 W at *T*_a_ = 50°C, i.e. > 6 × s.d. greater than the mean for the remaining cuckoos), and we excluded data from this individual from our RMR analyses and descriptive statistics. Minimum RMR values measured at *T*_a_ < 35°C, below the respective *T*_uc_s of each species, varied from 0.64 ± 0.20 W and 0.69 ± 0.13 W in cuckoos and rollers, respectively, to 1.19 ± 0.31 W in starlings (Table 1; mass-specific values are provided in supplementary information, Table S1). In rollers and starlings the *T*_uc_ (i.e., *T*_a_ inflection) ranged from 31°C in starlings to 47.5°C in rollers. No clear inflection point could be identified for cuckoos, and RMR increased across the entire range of *T*_a_ tested (Fig. 2). In all three species, RMR increased linearly and significantly at *T*_a_ > *T*_uc_ (cuckoo, t_1,23_=2.67, p<0.05; rollers, t_1,25_ = 2.59, p<0.05; starlings, t_1,29_=4.13, p<0.001) (Fig. 2). Maximum RMR observed at maximum *T*_a_ was higher in the starlings, with an average of 1.91 W, and lowest in the rollers at ~ 1.10 W. Maximum RMR was 0.96 W and 2.77 W in two cuckoo individuals, respectively (Table 1). The average magnitudes of RMR elevations above estimated thermoneutral levels (max. RMR / min. RMR) were generally below 1.6, with the exception of a single cuckoo that showed a 3-fold increase in RMR (Table 1). At the highest test *T*_a_ shared among the three species, RMR was quantitatively similar in cuckoos and rollers, but ~2-fold higher in the starlings (Table 1).

**Figure 2:**
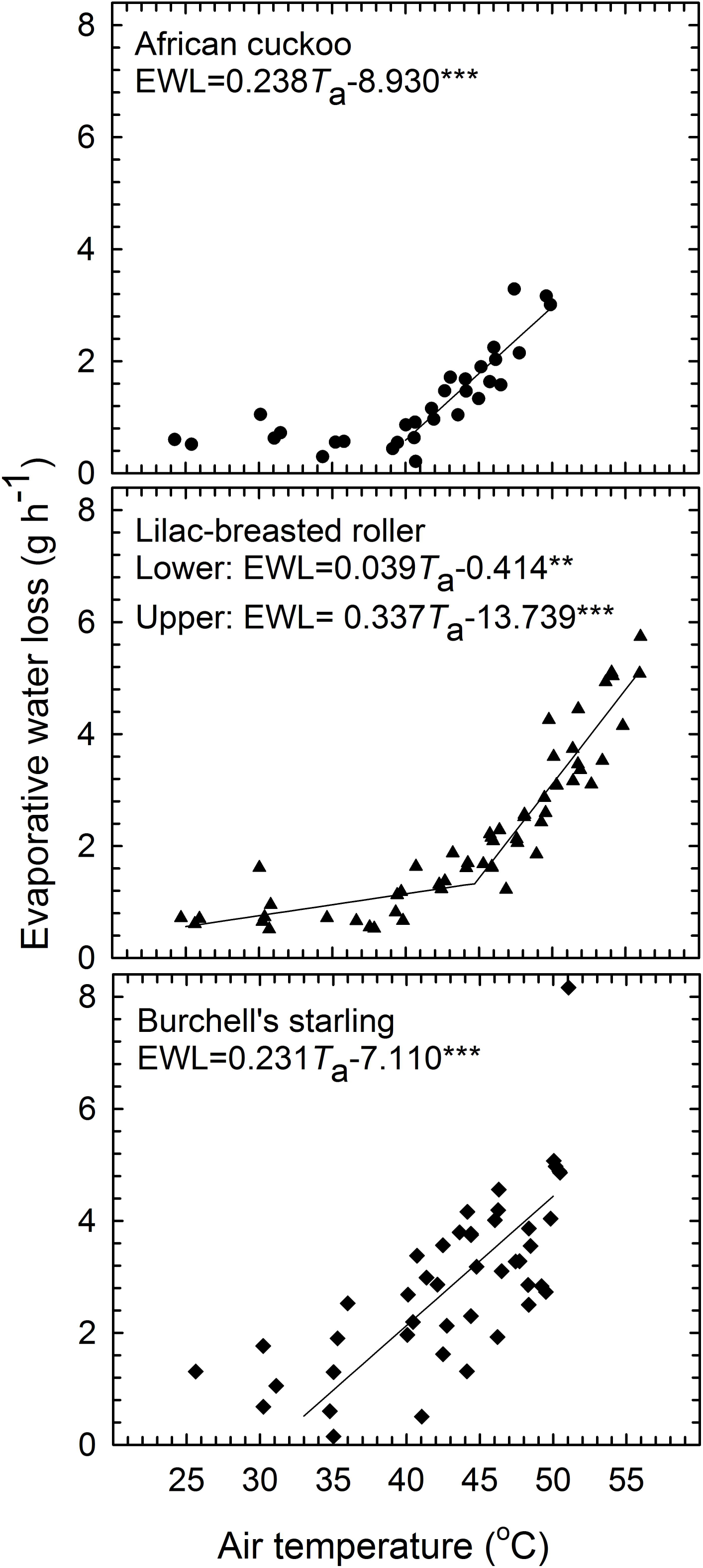
Resting metabolic rate (RMR) in African cuckoos (*Cuculus gularis*; n=5), lilac-breasted rollers (*Coracias caudatus*; n=10), and Burchell’s starlings (*Lamprotornis australis*; n=7) as a result of air temperature (*T*_a_). The trendlines represent relationships between *T*_b_ and *T*_a_ above an upper inflection *T*_a_ (see methods). The slopes and intercepts were calculated using linear mixed-effects models. Significant relationships are represented by asterisks: * = p < 0.05, ** = p < 0.01, *** = p < 0.001.

### Evaporative water loss

Minimum EWL values measured at *T*_a_s below the respective *T*_uc_ of each species ranged from 0.48 ± 0.14 g h^−1^ in cuckoos to 1.00 ± 0.67 g h^−1^ in starlings (Table 1; mass-specific values, TableS1). The inflection *T*_a_ varied by more than 10 °C among the three species; inflection *T*_a_ was 33.3 °C in starlings and greater than 42 °C in cuckoos and rollers (Table 1). Whereas cuckoos and starlings showed stable EWL rates below their respective inflection *T*_a_s, rollers showed a shallow but significant increase in EWL to 1.65 ± 0.35 g h^−1^ at *T*_a_ = 44.7°C (t_1,22_=3.22, p<0.01; Fig. 3). Above the inflection *T*_a_ EWL increased linearly and significantly in all three species (cuckoo, t_1,20_=13.22, p<0.001; rollers t_1,33_= 11.64, p<0.001; starlings, t_1,28_=11.133, p<0.001; Fig. 3). Maximum elevations in EWL were around 5-fold (>5.5-fold in cuckoos and rollers) above thermoneutral values in all three species (Fig. 3, Table 1). Whereas rollers showed maximum EHL/MHP ratios exceeding 3 at the maximum *T*_a_s tested, cuckoos and starlings generally showed values below 2 (Fig. 4). At the highest shared *T*_a_ cuckoos and rollers showed similar EWL rates and evaporative efficiencies (Table 1). In contrast, starlings showed EWL of ~ 3.40 g h^−1^, and a slightly lower efficiency compared to the former species (Table 1).

**Figure 3:**
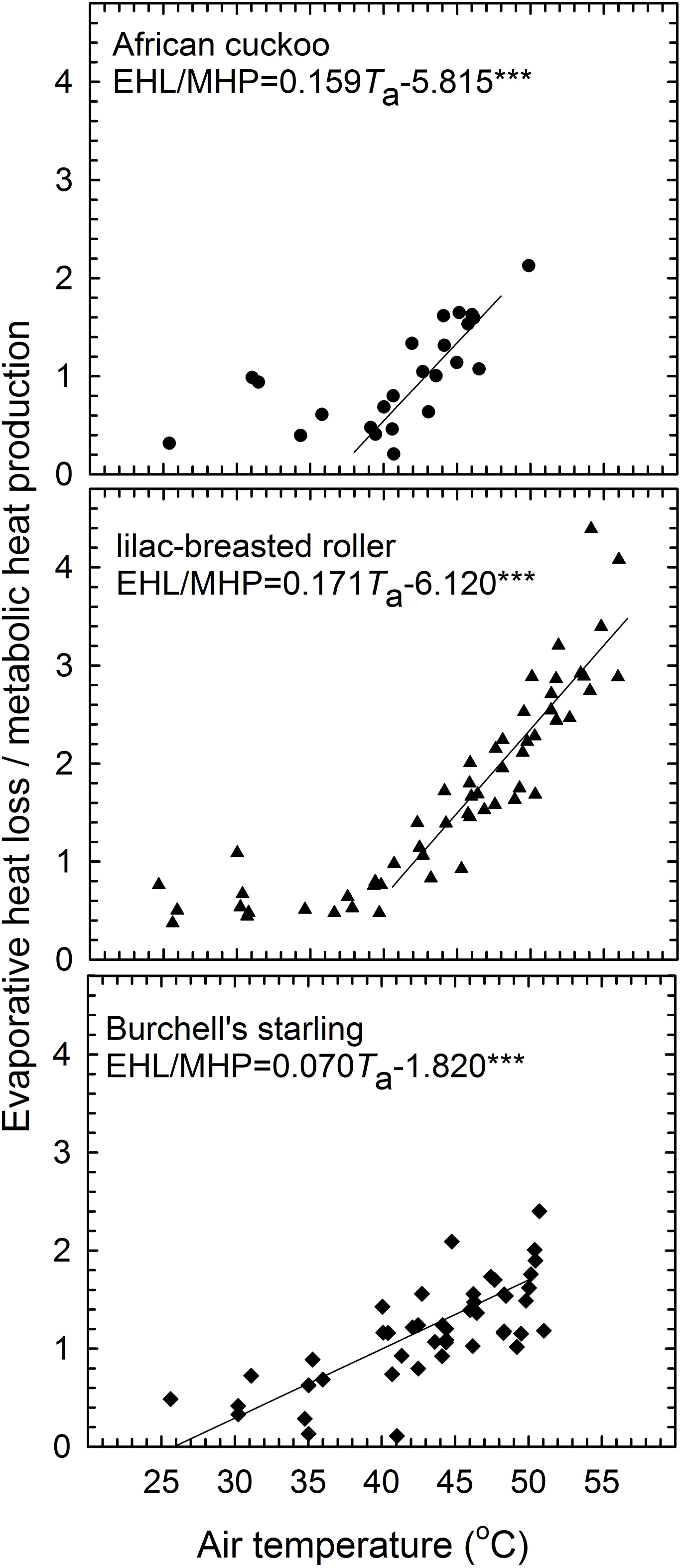
Evaporative water loss (EWL) in African cuckoos (*Cuculus gularis*; n=6), lilac-breasted rollers (*Coracias caudatus*; n=10), and Burchell’s starlings (*Lamprotornis australis*; n=7) as a result of air temperature (*T*_a_). The trendlines represent relationships between *T*_b_ and *T*_a_ above an upper inflection *T*_a_ (see methods). The slopes and intercepts were calculated using linear mixed-effects models. For lilac-breasted rollers two significant relationships are shown (25 °C < *T*_a_ < 44.7 °C; and *T*_a_ > 44.7 °C). Significant relationships are represented by asterisks: * = p < 0.05, ** = p < 0.01, *** = p < 0.001.

**Figure 4:**
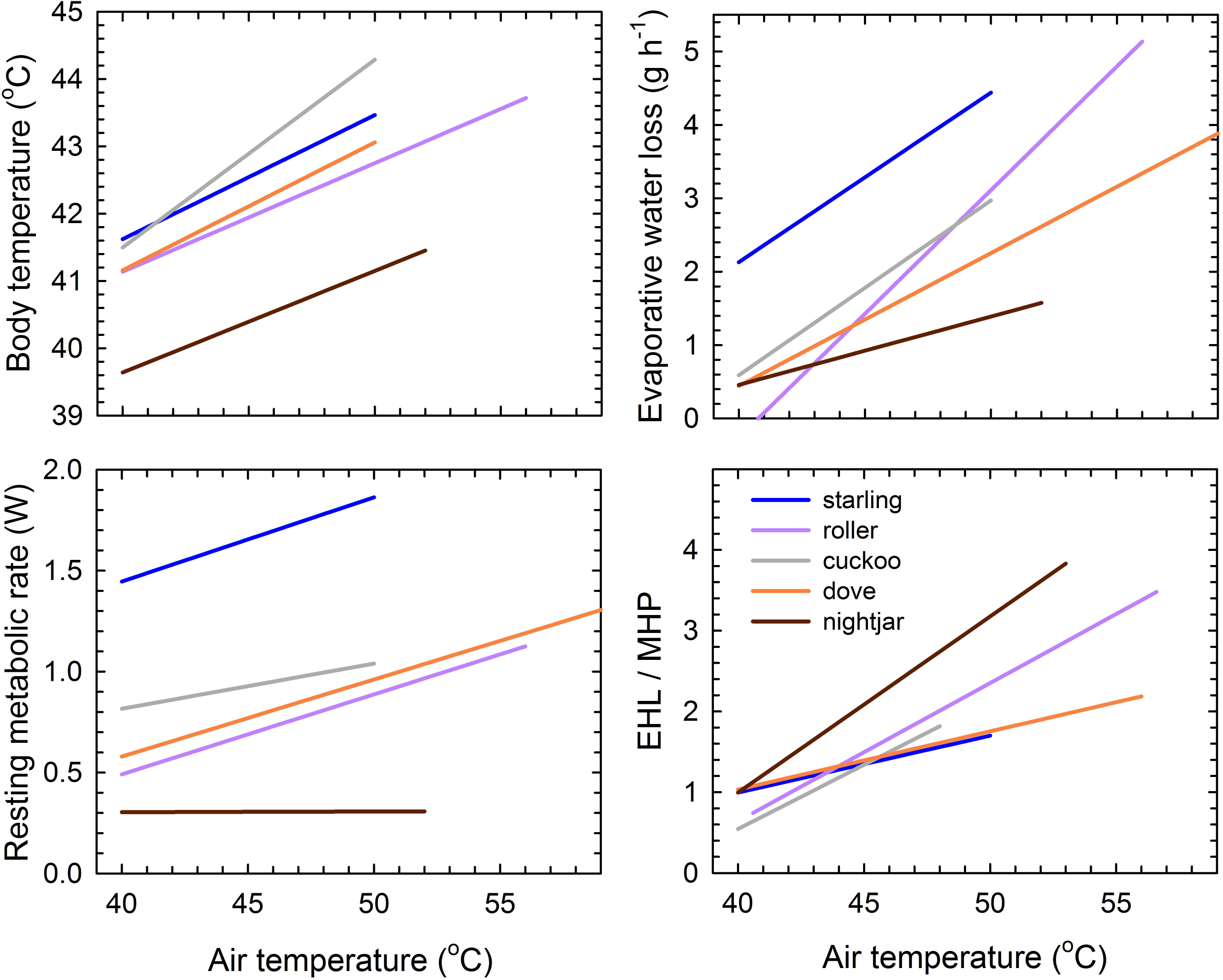
Ratio of evaporative heat loss (EHL) to metabolic heat production (MHP) in African cuckoos (*Cuculus gularis*; n=6), lilac-breasted rollers (*Coracias caudatus*; n=10), and Burchell’s starlings (*Lamprotornis australis*; n=7) as a result of air temperature (*T*_a_). The trendlines represent relationships between *T*_b_ and *T*_a_ above an upper inflection *T*_a_ (see methods). The slopes and intercepts were calculated using linear mixed-effects models. Significant relationships are represented by asterisks: * = p < 0.05, ** = p < 0.01, *** = p < 0.001.

### Variation in thermoregulation at high air temperatures

The *T*_b_ of cuckoos and doves increased more rapidly compared to rollers and nightjars (95% CL did not overlap, Fig. 5, Table 2). Starlings showed an intermediate slope of *T*_b_, with 95% CL overlapping with most other species. The slope of RMR at *T*_a_ above thermoneutrality was similar among species, with the exception of nightjars which showed almost no increase in RMR above 40°C and no overlap in 95% CL with any of the other species (Fig. 5, Table 2). The slope of EWL above the inflection *T*_a_ did not differ among cuckoos, starlings and doves, but was significantly steeper in rollers and significantly shallower in nightjars compared to all other species (Fig. 5, Table 2). The slopes of EHL/MHP were similar in doves and starlings, and significantly lower than in all the other species (Fig. 5; Table 2). Nightjars showed a steeper EHL/MHP slope compared to rollers and cuckoos, yet overlapped marginally with the 95% CL of these species.

**Figure 5:**
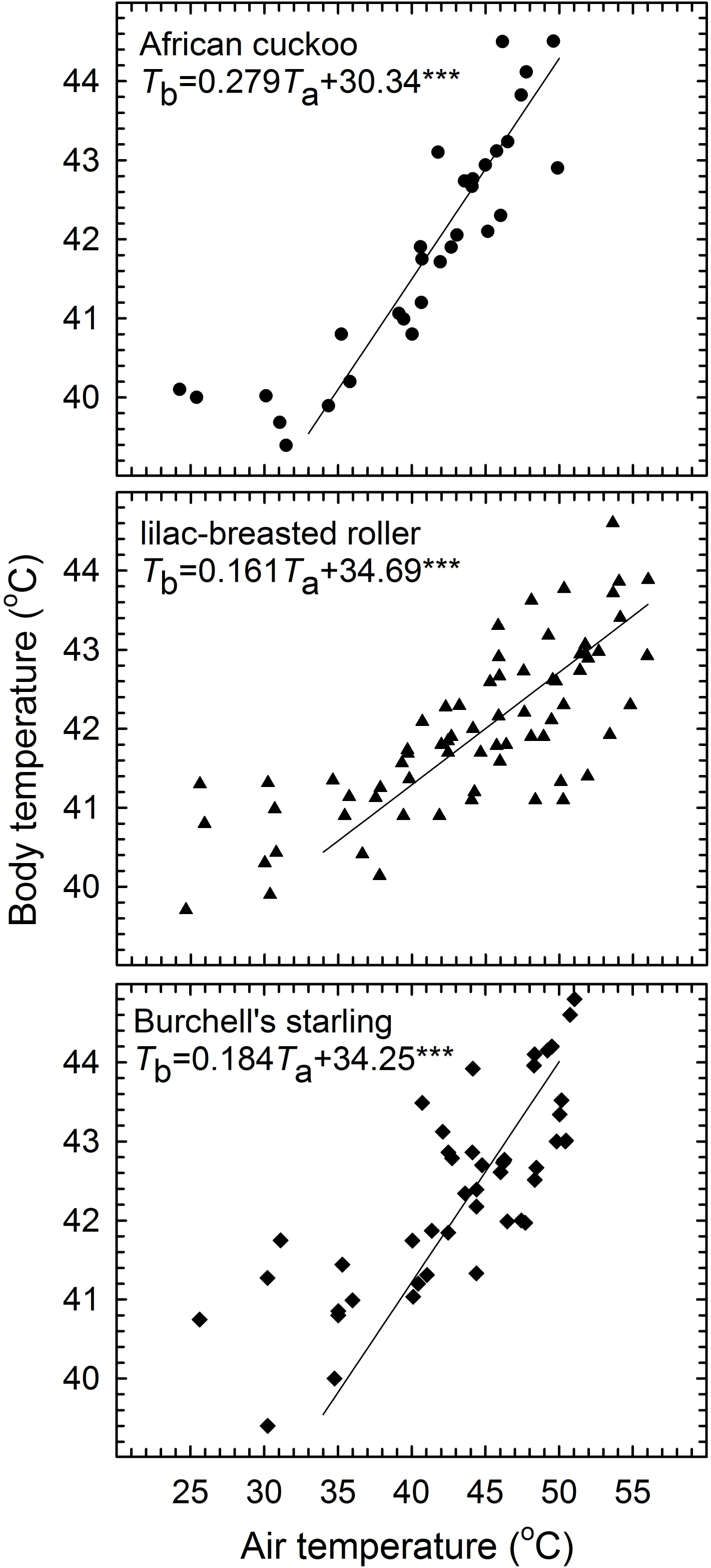
Physiological responses as a result of temperature in five desert species making use of varying modes of respiratory heat dissipation. These include African cuckoos (cuckoo; *Cuculus gularis*, this study) and laughing doves [dove; *Spilopelia senegalensis* (McKechnie et al. 2016a)] make use of gular flutter probably synchronised with panting. Lilac-breasted rollers (roller; *Coracias caudatus*, this study) and Burchell’s starlings (starling; *Lamprotornis australis*, this study) make use of panting only. Freckled nightjars [nightjar; *Caprimulgis tristigma* (O’Connor et al., 2017)] make use of rapid gular, but probably asynchronously to breathing rates. The slopes and 95% confidence levels of the relationship for each species are displayed in Table 2).

**Table 2:**
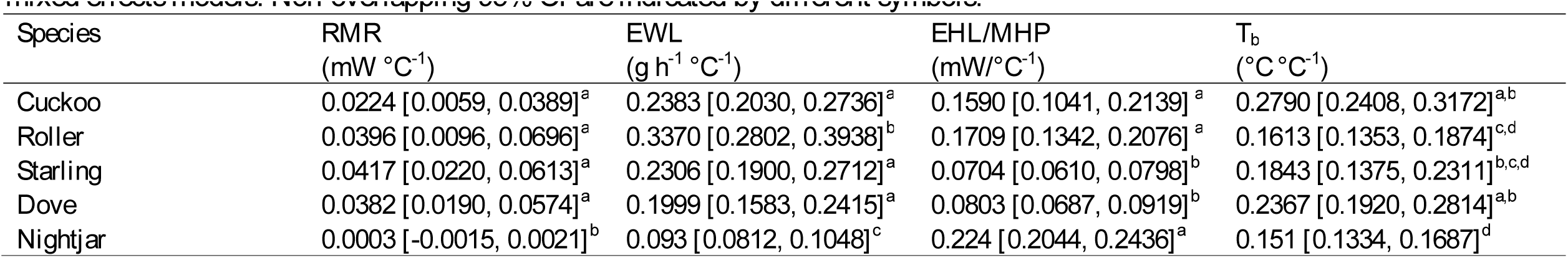
Slopes and upper and lower 95% confidence intervals (CI) for African cuckoos (cuckoo; *Cuculus gularis*, this study), Lilac-breasted rollers (roller; *Coracias caudatus*, this study), Burchell’s starlings (starling; *Lamprotornis australis*, this study), laughing doves [dove; *Spilopelia senegalensis* (McKechnie et al., 2016a)], and Freckled nightjars [nightjar; *Caprimulgis tristigma* (O’Connor et al., 2017)] derived from linear mixed effects models. Non-overlapping 95% CI are indicated by different symbols.

## Discussion

All three species studied investigated here showed elevations in *T*_b_, RMR and EWL at *T*_a_ > 40°C, qualitatively consistent with typical avian patterns. Despite their similarity in M_b_, the three study species showed substantial variation in patterns of *T*_b_, RMR and EWL at high *T*_a_. The comparatively shallow elevations in RMR in rollers were associated with the highest evaporative efficiency, and this species tolerated the highest *T*_a_. Rollers showed very small elevations in *T*_b_, associated with high evaporative efficiency. These patterns in rollers are unexpected since they appear to use panting as the primary EHL pathway. In contrast, at lower *T*_a_ both starlings and cuckoos showed sharp increases in *T*_b_ and RMR as well as lower evaporative efficiencies. Yet, the *T*_a_ at which these responses were initiated, patterns of *T*_b_ regulation and EWL elevations were strikingly different among these two species. Additionally, our five-order comparison reveals that all species, with the exception of caprimulgids (O’Connor et al., 2017), clearly showed elevated metabolic costs of heat dissipation (as defined by the slope of RMR above the inflection point) (Fig. 5). These findings illustrate how little we know about the phylogenetic variation in avian evaporative cooling pathways, and specifically the functional significance, selective pressures and evolution of these seemingly diverse pathways for heat tolerance.

### Body temperature and heat tolerance limits

The mean *T*_b_ of cuckoos at *T*_a_ below 35°C (39.8 ± 0.30°C) was 1.5°C lower than the mean active phase *T*_b_ values for Cuculiformes (41.3 °C), but the mean *T*_b_s of starlings and rollers were within 0.5°C of the mean values reported for Passeriformes and Coraciiformes, respectively (Prinzinger et al., 1991). In all three species, the inflection *T*_a_ for *T*_b_ was < 40°C (Fig. 1). These values are lower than the inflection *T*_a_s reported for columbids, quail, and smaller passerines (generally *T*_a_ ~ 40°C) (McKechnie et al., 2016a; Smith et al., 2015; Whitfield et al., 2015). Two species of Caprimulgiform also show a *T*_b_ inflection at *T*_a_ < 40°C (O’Connor et al., 2017).

In calm birds, maximum *T*_b_ was >44.5°C in a cuckoo at *T*_a_ = 49.6°C, 44.1 °C in a starling at *T*_a_ = 49.2°C and 44.6°C in a roller at *T*_a_ = 53°C (Fig. 1). The two cuckoos lost coordination at *T*_b_ > 44°C, suggesting this taxon may have lower critical thermal limits than Passeriformes and Columbiforms, where several species have been shown to tolerate *T*_b_ > 45°C (Dawson, 1954; McKechnie et al., 2016a; Smith et al., 2015; Smith et al., 2017; Whitfield et al., 2015). Several smaller passerines also reached thermal limits as *T_a_* approached 50°C (McKechnie et al., 2017; Smith et al., 2017; Whitfield et al., 2015) However, *T_a_* = 50°C is probably an underestimation of the heat tolerance limits of cuckoos and starlings, as we did not reach *T*_a_ at which calm individuals showed a plateau in EWL accompanied by an increasing *T*_b_ (Whitfield et al., 2015). In contrast to cuckoos and starlings, rollers defended *T*_b_ over a much larger *T*_a_-*T*_b_ gradient, tolerating *T*_a_ ≈ 56°C, with a mean *T*_b_ ≈ 43°C (Fig. 1). The *T*_b_ patterns of the coraciiform in our study at *T*_a_ approaching 60°C was therefore quantitatively similar to the thermoregulatory patterns observed in Columbiformes (McKechnie et al., 2016a; Smith et al., 2015).

### Resting metabolic rate

Individual variation in RMR tended to be greater in starlings and rollers compared to cuckoos. Some of this variation may be explained by the larger M_b_ ranges of starlings and rollers compared with cuckoos, but our data do not allow for a rigorous analyses of the scaling effects. Nevertheless, within species, individuals showed similar increasing trends in RMR elevations in slope estimates. The upper inflection *T*_a_ in RMR versus *T*_a_, i.e., the *T*_uc_ indicating increased energetic costs of respiratory EHL, generally occurs at *T*_a_ < 40°C (Calder and Schmidt-Nielsen, 1967; Dawson and Bennett, 1973; McNab and Bonaccorso, 1995; Tieleman and Williams, 2002; Whitfield et al., 2015). In all three species studied here, *T*_uc_ was not clearly related to increased efforts of respiratory EHL, unlike the case in Burchell’s sandgrouse (McKechnie et al., 2016b). Our data suggest that the initiation of gular flutter or panting was related to a threshold *T*_b_ of ~41.5°C, rather than a threshold *T*_a_ (Table 1). The cuckoos in our study did not show a clear upper inflection *T*_a_, contrasting with the marked *T*_uc_ associated with the onset of gular flutter in Burchell’s sandgrouse (*Pterocles burchelli*) (McKechnie et al., 2016b). At the *T*_a_ value at which we observed cuckoos initiating gular flutter (~42°C, Table 1), RMR had already increased by more than 50% above minimum resting levels. Similarly, in the starlings, the inflection *T*_a_ for RMR was well below the *T*_a_ at which this species initiated gaping behaviour, and lower than values in smaller passerines (Whitfield et al. 2015). The patterns in these two species are perhaps explained by a Q_10_ effect associated with *T*_b_ elevations, but our small sample sizes precluded us from drawing any firm conclusions; it is noteworthy that Weathers (1981) showed that, in several passerines, elevations in *T*_b_ do not affect RMR, but this idea has not been rigorously tested. In contrast to starlings and cuckoos, rollers maintained a low and stable RMR up to *T*_a_ = 47°C (Fig. 2), much higher than the *T*_a_ associated with gaping. This suggests that rollers kept metabolic heat loads low during the initial stages of panting. Our estimate of the upper *T*_uc_ in rollers is thus substantially higher than typically observed in passerines, which also make use of panting (McKechnie et al., 2017; Weathers, 1981; Whitfield et al., 2015).

Our data reveal considerable overlap in the slope of RMR increases above the *T*_uc_ in all three species studied here when compared to a similarly-sized dove (*S. senegalensis*) (Fig. 5, Table 2). The authors of at least one previous study noted that the metabolic costs in species that pant (e.g. village weavers, *Ploceus cucullatus*) are similar to those in species which gular fluttering (e.g. Chinese painted quails, *Excalfactoria chinensis*), and additionally cutaneous EHL [e.g. Inca doves, *Scardafella inca* (Lasiewski and Seymour, 1972)]. However, one striking difference among the species we compare here is that doves and rollers reached maximum RMR at higher *T*_a_ and maintained lower *T*_b_ compared to cuckoos and starlings. Fractional elevations in RMR at *T*_a_ = 50°C compared to thermoneutral values were as high as 60% in starlings, 25% in cuckoos, and ~10% in rollers and doves. These increases fall broadly between values previously reported for passerines [30-90%; (McKechnie et al., 2017; Whitfield et al., 2015) and columbids (~7% (McKechnie et al., 2016a) over a similar *T*_a_ range (30°C to 50°C). The lack of metabolic elevations at *T*_a_s above those where gular flutter was initiated in the nightjars (*T*_a_ ~ 40°C) contrasts strongly with all the above-mentioned species (Fig. 5). Metabolic rates at high temperature thus seem not related to the two modes (gular flutter versus panting) these phylogenetically diverse species employ.

### Evaporative heat loss

Patterns of EWL followed the typical avian pattern, where EWL is minimal at *T*_a_ around 25-35°C and increases approximately linearly at *T*_a_s approaching or exceeding 40°C. The variation in inflection *T*_a_ and slope of EWL above these inflections is difficult to explain, and suggests the capacity to dissipate heat evaporatively may vary greatly among taxa of similar size.

Maximum capacity for evaporative cooling, expressed as EHL/MHP, was qualitatively related to the maximum *T*_a_ reached by our three study species. On average, cuckoos and starlings evaporated less than 200% of metabolic heat production at the maximum *T*_a_ reached (Fig. 4) – similar to patterns observed in passerines (McKechnie et al., 2017; Smith et al., 2017; Whitfield et al., 2015). The comparatively high RMR of starlings likely constrained their evaporative cooling efficiency as they had to evaporate water almost twice as rapidly compared to the other species to dissipate 100% of their metabolic heat. This may have broad-scale implications for heat loss in the order Passeriformes, especially in light of a recent study demonstrating high basal metabolism in this order (Londoño et al., 2015). Cuckoos showed a slope of EHL/MHP versus *T*_a_ only marginally shallower than rollers, yet despite their low RMR at *T*_a_ < 35°C did not achieve a high evaporative efficiency (Fig. 4); even though a very calm cuckoo dissipated 200% of its metabolic heat load at *T*_a_ = 50°C, the *T*_b_ of this individual was already ~ 43°C.

Rollers showed a mean maximum EHL/MHP of 3.6 (> 4 in one individual) (Fig. 4), quantitatively similar to values reported in Columbiformes (2.3 - 4.7) (McKechnie et al., 2016a; Smith et al., 2015). In fact, at *T*_a_ > 50°C rollers evaporated a greater fraction of metabolic heat compared to medium-sized columbids (140-g *S. capicola*, and 90-g *S. senegalensis*) occupying the same Kalahari Desert habitat (McKechnie et al., 2016a). This result is surprising as panting is often argued to be a metabolically costly process compared to gular flutter (Bartholomew et al., 1962; Whitfield et al., 2015). We suspect the pronounced heat tolerance in rollers may be functionally linked to the long periods they spend perched in exposed sites while hunting (Herremans, 2005; Smit et al., 2016). We do not know whether rollers rely on cutaneous evaporation at high *T*_a_, although we propose that evaporative cooling may be enhanced by the large gape of rollers increasing the buccal surface area for EWL (personal observation); a combination of large gape [~15% of skin surface area, (Cowles and Dawson, 1951)] and high evaporative efficiency also is common to the caprimulgids, but it would be interesting to investigate the contribution of CEWL to total EWL in rollers. Our understanding of phylogenetic variation in the partitioning of EWL into cutaneous and respiratory pathways remains very poorly studied, and beside for a few studies focussing on Columbiformes and Passeriformes (McKechnie and Wolf, 2004; Ro and Williams, 2010; Tieleman and Williams, 2002; Wolf and Walsberg, 1996), there are limited empirical data on how increased reliance on CEWL at high *T*_a_ reduces the energetic demands for respiratory heat dissipation.

### Preliminary comparisons of phylogenetic variation in thermoregulation under hot conditions

Although our study was not designed to directly quantify gular flutter versus panting in our study species, our results, combined with those of (Lasiewski and Seymour, 1972), reveal that the mechanisms of respiratory heat dissipation are not clearly related to evaporative cooling capacity and tolerance of high *T*_a_. Even Columbiformes, thought to rely greatly on cutaneous EHL seem to suffer reductions in evaporative efficiency when they start incorporating gular flutter or panting mechanisms [e.g., *S. capicola*, and *S. senegalensis*, (McKechnie et al., 2016a)]. Calder and Schmidt-Nielson (Calder and Schmidt-Nielsen, 1967) suggested the process of gular flutter may differ among taxa and determine the metabolic costs of ventilating the buccal cavity. These costs may include processes where the frequency and amplitude of breathing cycles are in synchrony with gular flutter rates [e.g. Strigiformes, Columbiformes and Cuculiformes (Bartholomew et al., 1968)], or asynchronous where rapid gular flutter is accompanied by slow breathing rates [e.g. some Pelicaniformes and some Caprimulgiformes (Bartholomew et al., 1968; Lasiewski and Bartholomew, 1966; Lasiewski and Seymour, 1972)]. Bartholomew et al. (1968) further suggested that species maintaining gular flutter rates asynchronous to breathing rates, such as the poorwill (*Phalaenoptilus nuttalli*), achieve very high evaporative cooling efficiencies compared to species where breathing is completely synchronised with gular fluttering (probably no more costly than achieved through panting alone). To the best of our knowledge, the role these different modes of gular flutter play in the efficiency of EHL has not been further investigated.

The phylogenetic variation in modes of avian respiratory EHL pathways is interesting, and there is no clear explanation for the dichotomy between gular flutter and/or panting among avian taxa. The prevalence of gular flutter in phylogenetically older taxa, such as Paleognathae and most basal Neognathae, suggests it is a plesiomorphic trait. This may indicate that the function of the hyoid bone, which ventilates the gular area, was modified in some clades of the Neognathae (e.g. Coraciiformes and Passeriformes). Interestingly, Psitticiformes [parrots, closely related to Passeriformes, (Hackett et al., 2008) are the only avian taxon in which lingual flutter (movement of tongue) is known to supplement panting to ventilate the buccal area (Bucher, 1981; Bucher, 1985). However, to the best of our knowledge, no studies have examined whether lingual flutter differs in efficiency from panting alone.

In conclusion, substantial phylogenetic variation exists in avian heat tolerance and evaporative cooling capacity, an observation reiterated by our comparisons among representatives of five orders. This variation may have far-reaching implications for the ecology and evolution of birds that routinely have to maintain *T*_b_ below environmental temperature. The consequences of this variation for water balance and risk of over-heating during very hot weather are strikingly illustrated by the observation that, among the species considered here at *T*_a_ ~50°C, rates of EWL expressed in term of body mass loss vary ~2-fold; from an equivalent of 2.3% *M*_b_ h^−1^ in nightjars to 4.6% *M*_b_ h^−1^ in starlings. Moreover, at the same *T*_a_ the *T*_b_ of starlings is ~3°C higher than that of nightjars. Quantifying phylogenetic diversity in avian thermal physiology under hot conditions is critical for testing hypotheses regarding physiological adaptation, and predicting vulnerability to the higher *T*_a_ and more frequent and intense heat waves associated with anthropogenic climate change (Albright et al., 2017). For instance, the 140-fold differences in slopes of RMR and EWL with increasing Ta, (RMR 0.0003 mW °C^−1^ in nightjars to 0.0417 mW °C^−1^ in starlings) and 3.6-fold (EWL 0.093 g h^−1^ °C^−1^ in nightjars to 0.337 g h^−1^ °C^−1^ in rollers) suggests that, even within similarly-sized species, the consequences of the predicted 4 °C increase in extreme *T*_a_ maxima for the 21^st^ Century (IPCC, 2007; IPCC, 2011) will vary substantially. The current phylogenetic sampling of data on avian thermoregulatory responses to heat is poor, and a thorough review of the functional roles of these mechanisms as determinants of evaporative cooling efficiency will require sampling more avian taxa from currently unstudied orders.

## Acknowledgements

The Scholtz and de Bruin families (South Africa) allowed us to conduct this research on their properties. We also thank Michelle Thompson, Matthew Noakes, Ryan O’Connor, and Mateo Garcia for assistance in the field and laboratory.

## Author contributions

B.O.W., A.E.M. and B.S. designed the study. M.C.W., B.S., W.A.T. and B.O.W. collected data, B.S. and A.R.G. analysed the data. B.S. wrote the manuscript.

## Abbreviations

CEWL: cutaneous evaporative water loss
EHL: evaporative heat loss
EWL: evaporative water loss
*M*_b_: body mass
MHP: metabolic heat production
RMR: Resting metabolic rate
CL: confidence limit
*T*_a_: air temperature
*T*_b_: body temperature
V_CO_2__: carbon dioxide production
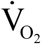: oxygen consumption

